# Model matters: Differential outcomes in traumatic optic neuropathy pathophysiology between blunt and blast-wave mediated head injuries

**DOI:** 10.1101/2023.05.25.542261

**Authors:** S.M. Hetzer, C. O’Connell, V. Lallo, M. Robson, N.K. Evanson

## Abstract

Over 3 million people in the United States live with long-term disability as a result of a traumatic brain injury (TBI). The purpose of this study was to characterize and compare two different animal models of TBI (blunt head trauma and blast TBI) to determine common and divergent characteristics of these models. With recent literature reviews noting the prevalence of visual system injury in animal models of TBI, coupled with clinical estimates of 50-75% of all TBI cases, we decided to assess commonalities, if they existed, through visual system injury. Blast, repeat blast, and blunt injury were induced in adult male mice to observe and quantify visual deficits. Retinal ganglion cell loss and axonal degeneration in the optic tract, superior colliculus, and lateral geniculate nuclei were examined to trace injury outcomes throughout major vision-associated areas. Optokinetic response, immunohistochemistry, and western blots were analyzed. Where a single blunt injury produces significant visual deficits a single blast injury appears to have less severe visual consequences. Visual deficits after repeat blasts are similar to a single blast. Single blast injury induces contralateral damage to right optic chiasm and tract whereas bilateral injury follows a single blunt injury. Repeat blast injuries are required to see degeneration patterns in downstream regions similar to the damage seen in a single blunt injury. This finding is further supported by Amyloid Precursor Protein (APP) staining in injured cohorts. Blunt injured groups present with staining 1.2 mm of the optic nerve, indicating axonal breakage closer to the optic chiasm. In blast groups, APP was identifiable in a bilateral pattern only in the geniculate nucleus. Evidence for unilateral neuronal degeneration in brain tissue with bilateral axonal ruptures are pivotal discoveries in this model differentiation. Analysis of the two injury models suggest there is a significant difference in the histological outcomes dependent on injury type, though visual system injury is likely present in more cases than are currently diagnosed clinically.

## Introduction

Traumatic brain injuries (TBIs) comprise 1/3^rd^ of all injury related deaths in the United States (Prevention, 2022). Occurring through a litany of mechanisms including improvised explosive devices, sports injuries, collisions, or falls, and ranging from mild to severe, millions are treated in the emergency department (ED) each year. Mild to moderate severity accounts for up 70-90% of ED visits for TBI, but this number is likely underreported as it only reflects those who seek hospital care(Daniel Laskowitz and Grant, 2016); and, even if mild TBIs do not incite noticeable deficits, we now know that there are chronic changes to white matter tracts in the brain termed traumatic/diffuse axonal injury (Johnson et al., 2013; Vieira et al., 2016). One white matter tract that is particularly vulnerable is the optic nerve, which connects retinal ganglion cells (RGCs) in the eye to the brain. Although damage to the axons of RGCs is not always the cause of TBI-induced visual deficits, 50-60% of TBI patients across the spectrum of severities and modalities of injury develop some type of visual impairment (Ventura et al., 2014). When injury is specific to the optic nerve, diagnosis is complicated because limitations of *in vivo* imaging, confounds of intraocular pressure, or a lack of visible damage obfuscate clinicians’ ability to diagnose traumatic optic neuropathy (TON) (Singman et al., 2016). Moreover, some patients do not report visual disturbance immediately after injury, and small changes in vision often go undiagnosed by physicians treating the TBI (Chan et al., 2019). This prevents us from completely understanding how prevalent TON is among TBI patients.

Study of optic nerve damage has relied on methods such as optic nerve crush, nerve transection, or ocular blast models, but these injury modalities are removed from the context of TBI. This TBI “context” is further complicated in laboratory settings because several methods of experimental TBI have been developed to model different types of injury (e.g., controlled cortical impact, fluid percussion, blunt head injury, and blast). Each has its own benefits and limitations for assessing aspects including regional-specific injury, spreading injuries, white matter injury, concussion, or explosive pressurized injuries. But which model is best? Which model accurately recapitulates human TBI? If human TBIs are unique to individuals, how can we use animal models that also isolate unique TBI characteristics to examine more *generalizable* consequences of injury? Over the last decade scientists have been asking these questions and are starting to approach them by formulating common data elements thereby establishing a uniform set of vocabulary for describing TBI outcomes (Smith et al., 2015). While this might have aided clinical diagnosis and categorization of TBI patients, there remains considerable variability in the basic science literature. At present, one of the only categorically agreed upon factors has been classification of injuries as open or closed head (Smith et al., 2015).

Exploring the means by which TBI induces generalizable/overlapping injury pathophysiology (i.e., outcomes associated with secondary injury, such as neuroinflammation, phagocytosis, axonal shearing/degeneration) is one path to understanding similarities among models. If we can find commonalities arising from any type of TBI, we might be able to provide better treatment guidelines or new ways of assessing deleterious outcomes. Our lab has induced traumatic axonal injury to the optic nerve using a closed-head weight drop TBI model in mice, and other similar models have reported parallel findings of injury in the optic tracts (Bashir et al., 2020; Bernardo-Colon et al., 2019; Evanson et al., 2018; Heldt et al., 2014; Tao et al., 2017; Wang et al., 2013). However, to our knowledge, direct comparisons between animal models of TBI to determine whether TON might be an overlapping phenotype have yet been performed. This study aimed to compare two commonly utilized preclinical models of closed-head mild TBI, blunt- and blast-mediated, with the intention of identifying convergent or modality-specific features within the visual system. Both blunt and blast TBI share injury pathology but are induced by distinct mechanisms (contusion versus acceleration/deceleration). As such, the elucidation of shared or unique mechanisms of injury to the visual system may lead to novel treatment approaches to attenuate injury progression in one or both TBI modalities.

## Methods

### Animals

Experiments were performed in 8-week-old male C57BL/6J mice (Jackson Laboratories, Bar Harbor, ME) and 8–9-week-old C57BL/6 Charles Rivers mice. Analyses for strain differences showed no effects of strain, so data were combined. Mice were housed under a 14h:10h light:dark schedule in pressurized individually ventilated cage racks, with 4 mice per cage, and were given ad libitum access to water and standard rodent chow. Animals habituated to the vivarium for at least one week prior to undergoing traumatic brain injury and subsequent procedures. A timeline is provided in Figure 1a. The University of Cincinnati Institutional Animal Care and Use Committee approved all experimental procedures (#s 23-02-09-02 and 20-11-05-01).

**Figure 1.**
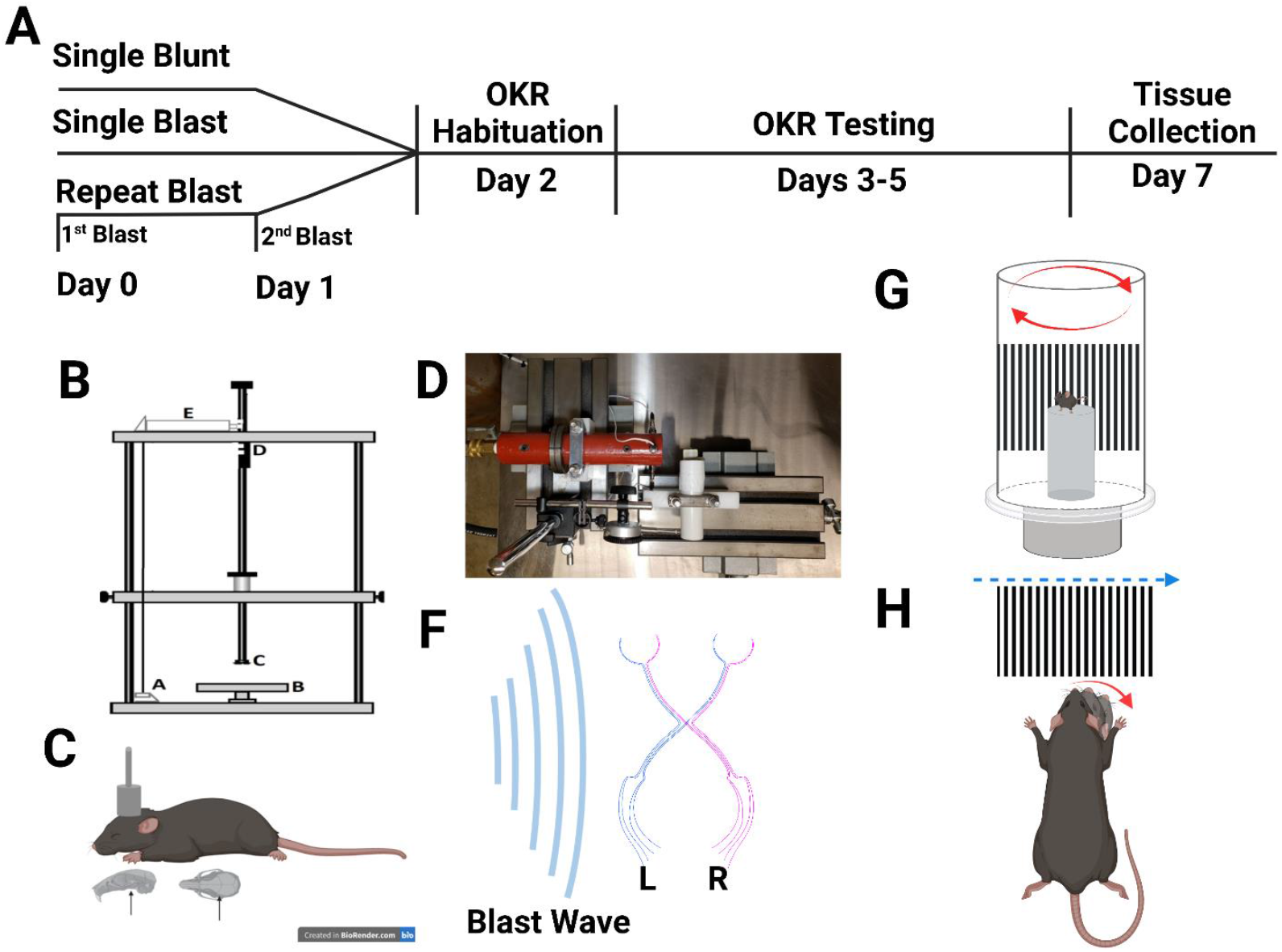
Overview of methods. (A) The experimental timeline for procedures is shown with 4 groups of mice (single blunt, single blast, repeat blast, and sham) tested for optokinetic nystagmus on days 3 -5 after TBI with tissue collection at day 7. (B) Shows an image of the blunt TBI weight-drop device and (C) a rough localization of injury when the weight is dropped. (D) shows the blast device used with (F) an illustration of the direction of the blast wave (blue lines) to show that the wave hits the left side of the head along the length of the mouse’s skull [L=left], [R=right]. (G) Shows a depiction of the optokinetic machine used for visual testing with (H) representative images of contrast grating and behavioral feature of the mouse’s head following the drum. OKR = Optokinetic Response

### Traumatic Brain Injury

#### Blunt Injury

Closed-head injury was performed by weight drop, as previously described (Cansler et al., 2020). Briefly, mice were anesthetized using isoflurane (2-3%), placed in prone position under a metal rod raised above the intact, unshaven scalp. The rod was dropped roughly above bregma (Fig. 1 b, c). Mice were observed for righting reflex time (RRT) before being returned to their home cages. Sham animals were anesthetized, weighed, and allowed to recover before being returned to their cages.

#### Blast Injury

Following a 30-minute acclimation period to the testing facility, subjects were anesthetized with 4% volume-to-volume isoflurane (VetEquip). Subjects were induced (Plane III, paralysis of intercostal muscles) and immediately subjected to blast TBI or sham treatments as previously described.(Logsdon et al., 2020) A second group of mice were subjected to a second blast 24 hours after the first (rblast). Briefly, subjects were placed within a polyvinylchloride (PVC) blast shield, perpendicular to blast wave front, shielding internal organs from injury, but leaving the head free to move. The blast device consists of a 2-piece machined steel shock tube apparatus, driven by compressed helium gas (Airgas). The driver and driven sections of the apparatus were separated by a 0.003” Mylar^®^ membrane (ePlastics; Fig. 1d, f) (Logsdon et al., 2020; Turner et al., 2013). Shock waves generated were scaled in intensity (≈ 1250 kPa) and duration (≈1 ms overpressure) for murine subjects.(Logsdon et al., 2020) Sham subjects were exposed to anesthesia, shielding tube, and noise, without exposure to shock wave. Immediately following TBI or sham treatments, subjects were immediately removed, and RRT was measured and recorded prior to subjects being returned to their respective home cages.

### Optokinetic Nystagmus Behavioral Testing

As described previously (Hetzer et al., 2021a), we tested visual and oculomotor function by measuring the involuntary optokinetic nystagmus response (OKR). Briefly, mice are placed on an immobile platform surrounded by a rotating Plexiglas drum (Fig. 1g). An exchangeable series of sine-wave gratings (black and white bars calibrated to specific cycles-per-degree (cpd) specifications; Fig. 1h) are placed around the inside of the drum. For these experiments, three gratings were used (0.26, 0.32, and 0.39 cpd) to examine a range of visual acuities with higher cpd gratings requiring greater visual acuity to respond to. Once in the drum, mice are left for a one-minute habituation period before the machine is turned on. The drum rotates at two revolutions per minute (rpm) for two minutes in both clockwise or counterclockwise rotation, followed by 30 seconds rest, and two minutes rotation in the opposite direction. Mice are exposed to one grating per day in random orders. Experimenters are blinded to conditions during both behavior and scoring. Videos are scored by trained experimenters and the number of OKRs is tallied. Two scorers review each video, and their tallies are averaged and assessed for appropriate inter-rater reliability.

### Histology

#### Tissue Collection

For immunohistochemical and immunofluorescence analyses, mice were euthanized using pentobarbital overdose seven days after TBI (timeline Fig. 1a). Mice were perfused transcardially with 4% paraformaldehyde in 0.02M phosphate-buffered saline (PBS) solution (pH 7.4). Brains were removed and post-fixed in 4% paraformaldehyde in 0.02M PBS overnight at 4°C, rinsed in 0.01M PBS, and immersed in 30% sucrose solution at 4°C until sectioning. Sucrose-saturated brains were frozen on dry ice and sectioned at 30μm using a sliding microtome (Leica, Bannockburn, IL). Sections were stored in cryoprotectant solution (0.01M PBS, polyvinyl-pyrrolidone (Sigma Aldrich, Cat: PVP-40), ethylene glycol (ThermoFisher, Cat : E178-4), and sucrose (ThermoFisher, Cat: S5-3) at -20°C until staining was performed. In a sma **l** cohort, sham and blunt TBI mice were perfused, and only optic nerves were collected for whole mount staining described below. For western blotting, both eyes were taken after injection with pentobarbital once no toe pinch or eye blink responses were detected and before perfusion. After removal, eyes were placed in ice cold 0.01M PBS, and the retina was removed within 5 minutes, immersed in cold lysis buffer (described below), and frozen on dry ice. Retinas were stored at - 80°C until used for western blotting.

#### Amyloid Precursor Protein

Free-floating brain tissue was stained using mouse monoclonal antibodies against Amyloid Precursor Protein (ThermoFisher, CAT: 22C11; RRID:AB_2893281; 1:1000 dilution) overnight at 4°C. The next day, tissue was washed, incubated for 1 hour at room temperature in biotinylated anti-mouse secondary antibody at 1:500 dilution (Vector Laboratories; cat # BA-9200; RRID: AB_2336171), followed by incubation in ABC Media 1:800 (Vector Laboratories; cat # PK-6100; RRID: AB_ 2336817) for 60 minutes at room temperature, and about 8-10 minutes exposure to DAB (3,3′-Diaminobenzidine; Sigma-Aldrich D5637). Slides were coverslipped using DPX Mounting Media (Sigma-Aldrich, CAT: 06522).

#### FluoroJade B

Fluoro-Jade B (FJ-B; Histo-Chem, Jackson, AR; CAT# 1FJB), a marker for degenerating neurons and axons,(Schmued and Hopkins, 2000) was used to stain tissue sections according to the manufacturer’s directions with slight modifications to avoid high background. Sections were incubated in 0.06% potassium permanganate for 10 minutes, rather than 15min, then submerged in 0.0004% FJ-B solution for 20 minutes. After staining, slides air dried in the dark. Slides were stored without coverslips in a slide box and imaged immediately, again, to reduce background and quenching of positive signals.

#### IBA-1 & CD68

Free-floating brain tissue was stained for microglia and for a microglia-specific phagocytic marker via staining with rabbit monoclonal IBA-1 (Ionized calcium binding adaptor molecule 1; Wako, Cat: 019-19741, RRID:AB_839504) 1:1000 and rat monoclonal CD-68 (Cluster of Differentiation 68; Bio-Rad, MCA1957, RRID:AB_322219) 1:750 overnight at 4°C. Tissue was then incubated overnight again in a cocktail of donkey anti-rabbit Cy3 (1:1000; ThermoFisher Invitrogen, CAT: A10520, RRID:AB_2534029) and goat anti-rat 647 (1:1000; ThermoFisher Invitrogen, Cat: A21247, RRID:AB_141778). Slides were coverslipped using the antifading polyvinyl alcohol mounting medium (Sigma-Aldrich, Cat: 10981; St. Louis, MO).

### Image Analysis

For image analysis both left and right sides of various optic system regions of interest (ROIs) were examined including the Optic Nerve, Optic Chiasm, Optic Tract (OT), Lateral Geniculate Nucleus (LGN), Superior Colliculi (SC), and the accessory optic system’s dorsal terminal nucleus (DTN). For APP images, high resolution, 60x large image composites were stitched together using a Nikon C2 Plus Confocal Microscope (Nikon Corporation, Melville, New York). These images were qualitatively examined for APP staining in the optic nerve, OT, LGN, and SC. For FJB analysis, images were taken at 20x on the same Nikon microscope. FJB positive area was measured using Nikon Elements Analysis software by thresholding the image to count the area of positive FJ stain within a bezoar ROI of areas of interest only (Nikon, Melville, NY). IBA-1/CD68 images were taken at 40x for both channels (i.e., wavelength) and included a merged image. IBA-1/CD68 analysis was completed using Image J to threshold positive CD68 area and the cell counting feature to count the number of microglia somata. All images were taken and analysed by a blinded researcher.

### Western Blotting

Retinas were homogenized in RIPA lysis buffer (Sigma-Aldrich, Cat: R0278) using a pellet homogenizer, centrifuged at 3000 rpm for 20 minutes, and supernatant removed for protein concentration analysis using a BCA protein assay (Pierce BCA Protein Assay Kit; ThermoFisher Scientific; cat # 23227). Twenty μg samples were loaded into SDS-PAGE gels and transferred onto Amersham Hybond-P 0.45μm PVDF membranes (GE Life Sciences, Pittsburgh, PA; CAT: GE 10600029). Membranes were incubated in Fisher No-Stain Total Protein Stain (ThermoFisher, CAT: A44449) as per manufacturer instructions, blocked in 5% non-fat milk for 1 hour at room temperature, then incubated overnight in polyclonal anti-RBPMS (RNA Binding Protein Multiple Partners; 1:1000; ThermoFisher, CAT: PA5-31231) at 4ºC. Membranes were rinsed in TBST then incubated in anti-rabbit HRP for 1.5-2 hours at room temperature. Blots were imaged using an iBright™ CL750 Imaging System (ThermoFisher, Waltham, MA) and analyzed using Image J. Some blots were stripped following imaging in stripping buffer (β-mercaptoethanol, 20% Sodium Dodecyl Sulfate, and 1M Tris-HCl pH 6.8) for 30 minutes at 50ºC, washed, re-blocked, and exposed to the same immunoblotting steps as above. Due to the large number of subjects, multiple blots were needed, so a constant inter-membrane control (homogenized naive mouse brain) was used on each blot and samples were first normalized to total protein then to the intermembrane control. Band contrast analyses were completed using ImageJ.

### Statistical Analyses

OKR data for overall differences among groups was done by one-way ANOVA within each spatial frequency separately (a 2-way RM ANOVA including difference between grating was not performed because this difference is expected, as we have previously discussed).(Hetzer et al., 2021a) For further analysis of potential left and right eye differences, OKR was analyzed by 2-way ANOVA (injury x spin direction). Post Hoc analysis was done with the Holm-Sidak test for multiple comparisons. FJB and IBA1/CD68 data were analyzed by 2-way RM ANOVA (injury x left/right). Western Blots were analyzed using one-way ANOVA since the left and right retina of blunt mice was combined. All analyses were computed and graphed using GraphPad Prism 9 (GraphPad Software, San Diego, California).

## Results

### Injury location is different between TBI groups

APP staining revealed clear differences among comparison groups. Beginning by staining brain tissue, we saw no positive staining in sham tissue or blunt injured mice (see supplementary image files). Blast and repeat blast mice, on the other hand, displayed robust staining in both the OT (Fig. 2a-d) and LGN (Fig. 3a-d) in both left and right sides, and with subjectively more staining on the right (contralateral) side of the brain.

**Figure 2.**
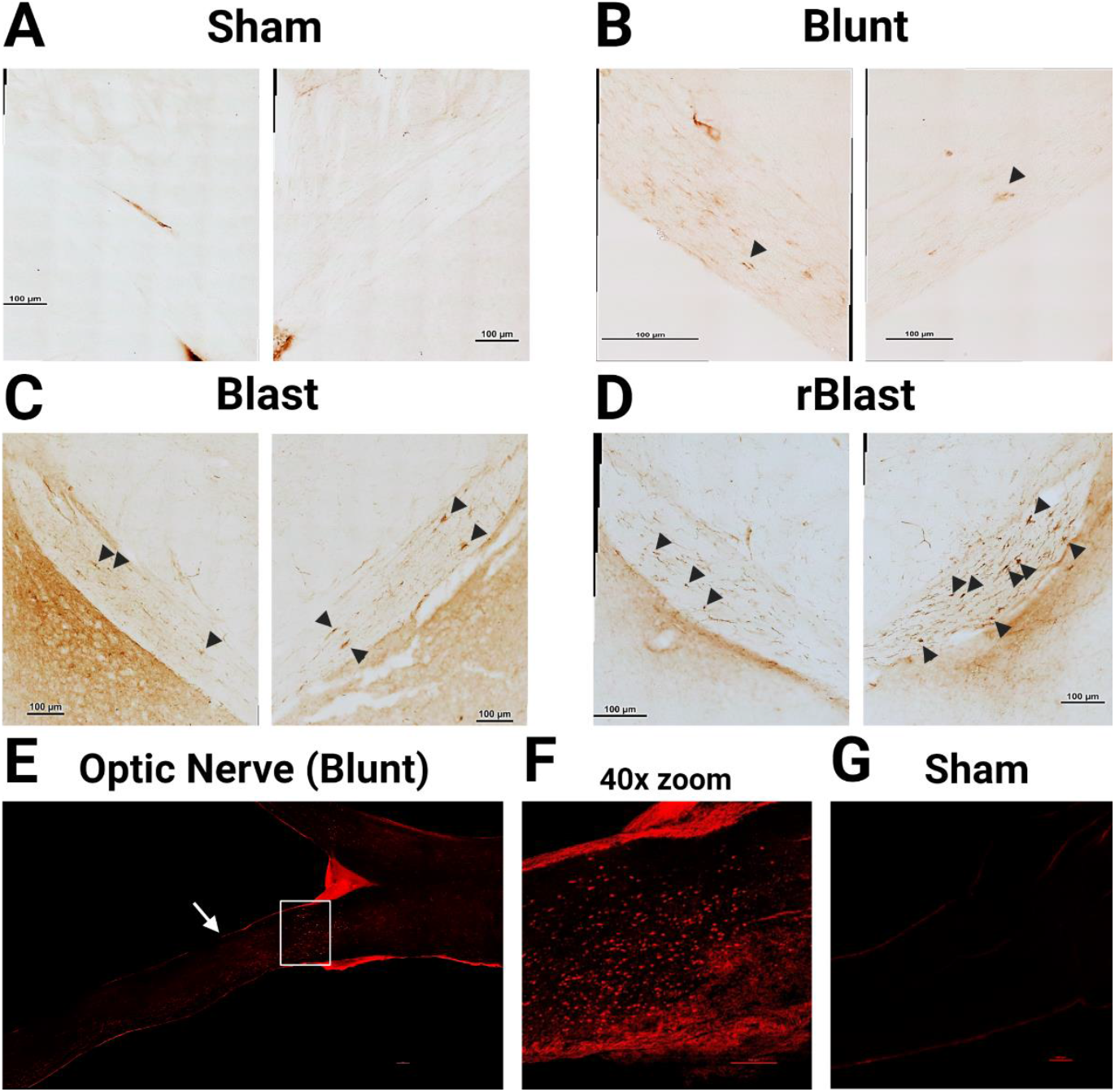
Estimated location of axon breakage. Using APP staining to find where axon breakage occurred, qualitative assessment of DAB bright field images shows no APP in (A) sham optic tracts and (B) minimal to no APP in blunt injured mice (black arrows). (C) Increasing levels of APP are found in OTs of blast mice bilaterally with more APP given (D) repeated blast injures. (E) Separate fluorescent immunostaining for APP in blunt injured optic nerves revealed the injury location within 1.2 mm proximal to the chiasm (white arrow), (F) 40x zoon of white box in E. (G) Shows no APP in sham mice. Bright field images consist of a large image stitch of 60x magnification. (E) is a stitched 10x image, and (G) is a stitched 20x image. Scale bars represent 100 μm. APP = Amyloid Precursor Protein, OT = optic tract, rBlast = repeat blast

**Figure 3.**
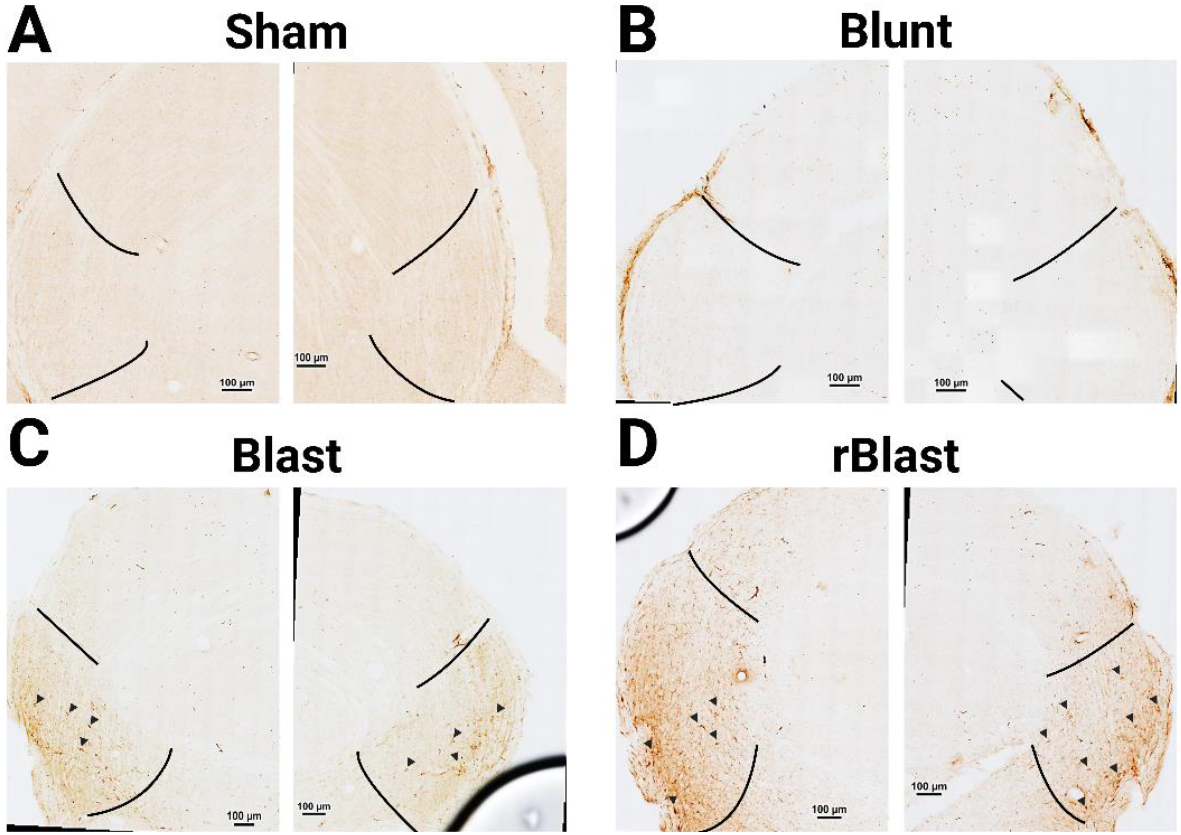
Blast Injury induces differential thalamic injury. APP-stained tissue was also imaged in the optic tract projection target, the lateral geniculate nucleus. The LGN is comprised of three divisions, the dorsal (topmost area), intergeniculate leaflet (covered by the topmost black lines), and ventral nuclei (bottom most area inside the two back lines) which each project to different downstream optic regions. There was no positive staining in (A) sham or (B) blunt injured LGNs. (C) single blast TBI mice show positive staining in only the ventral LGN (black arrows) while (D) repeat blast seems to lead to more severe APP staining that is present in both dorsal and ventral regions. Images are comprised of stitched 60x images, Scale bars represent 100μm. APP = amyloid precursor protein, LGN= lateral geniculate nucleus, rBlast = repeat blast injury.

The neurodegenerative stain, FJ-B, revealed a different set of findings, however. There was no positive staining in control mice, but blunt injured mice presented with significant levels of FJB bilaterally in the optic chiasm (p<0.001; Fig. 4a, b), OT (p<0.001; Fig. 4c, d), LGN (p<0.004; Fig.5a-c), SC (p<0.03; Fig. 5d-f), and DTN (p<0.003; Sup. Fig. 1a-c) versus sham. In the chiasm, blunt injured mice had significantly more FJB than rblast mice (p=0.005) but similar levels to single blast in the right OT (p=0.7). Blast injured mice only showed positive staining in the right chiasm (perfectly bisected through the middle; right only p=0.03; Fig. 4b) and OT (p=0.03; Fig. 4c, d) with little to no positive staining in the LGN (p>0.9), SC (p=0.9), or DTN (p=0.9). Repeatedly injured blast mice also predominantly presented with contralateral FJB that was significantly elevated in the right chiasm (p=0.01; Fig. 4a, b), OT (p=0.005; Fig. 4c, d) and SC (p=0.001; Fig. 5d-f). Although positive staining was present, and comparable to blunt mice in a few rblast mice, there was not a significant elevation in the LGN (p=0.4; Fig. 5a-c) or DTN (p=0.9; Sup. Fig. 1a-c). There was, however, a significant increase in FJB between left and right DTN within rblast mice (p=0.01). Because we saw FJB but no APP in blunt injured mouse brain tissue, we collected whole optic nerves for APP staining in blunt injured mice and sham animals. We found that the location of axon shearing in the blunt model is outside the brain in the 1.2mm of the optic nerve proximal to the optic chiasm (Fig. 2e-g).

**Figure 4.**
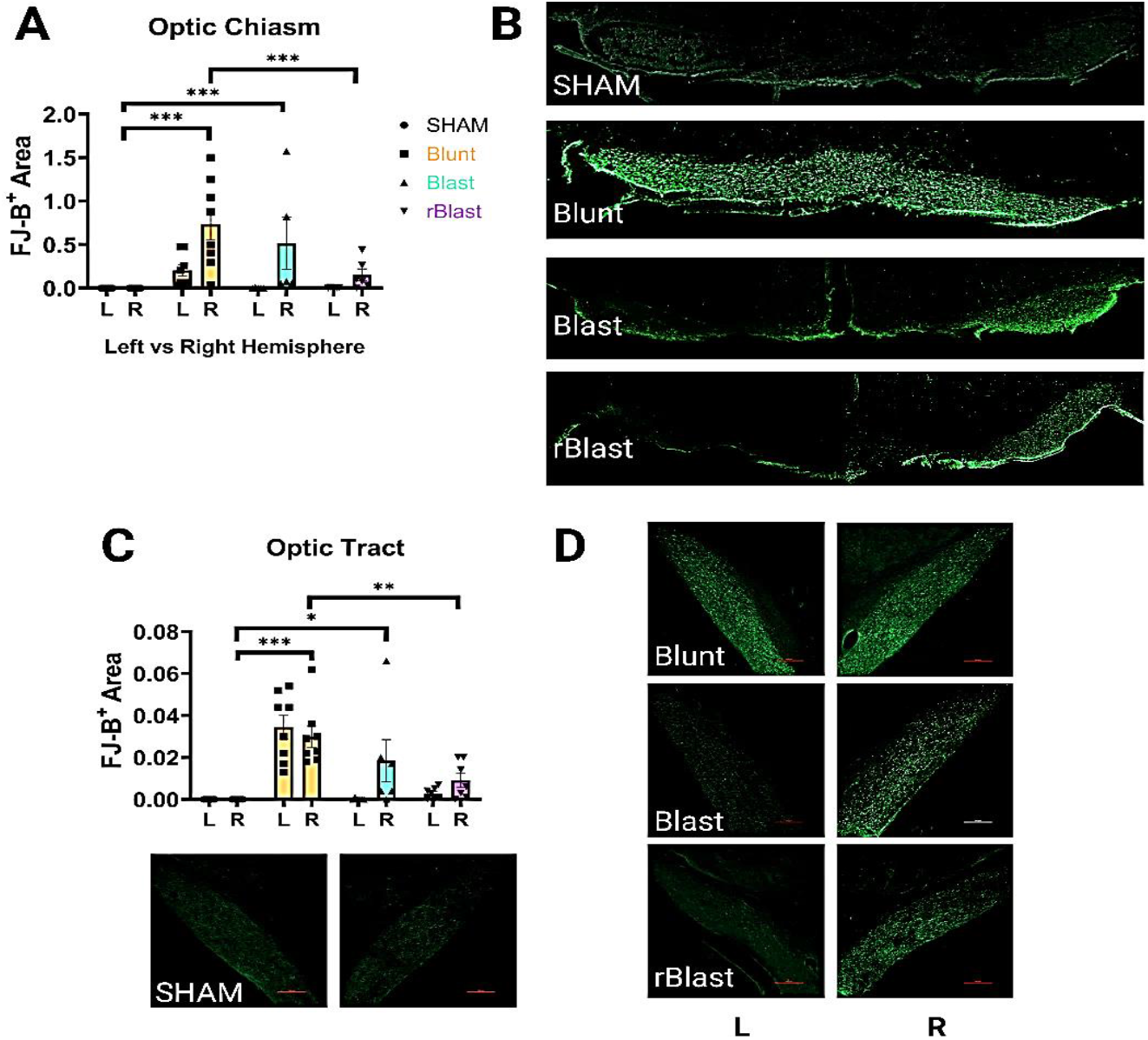
Optic tract axonal degeneration is bilateral in blunt mice but unilateral in blast mice. (A) Significant increases in FluoroJade-B positive area were found in all injury groups’ optic chiasms compared to sham, but this increase was only significantly present in the right OX of blast mice. (B) 10x representative images of optic chiasms. (C) The optic tract showed a similar pattern of degeneration. (D) 20x representative images of optic tracts. L= left, R=right, * p<0.01, ** p<0.001, ***P<0.0001, scale bars represent 100μm. FJ-B = FluoroJade-B, L = left, R = Right, rBlast = repeat blast TBI

**Figure 5.**
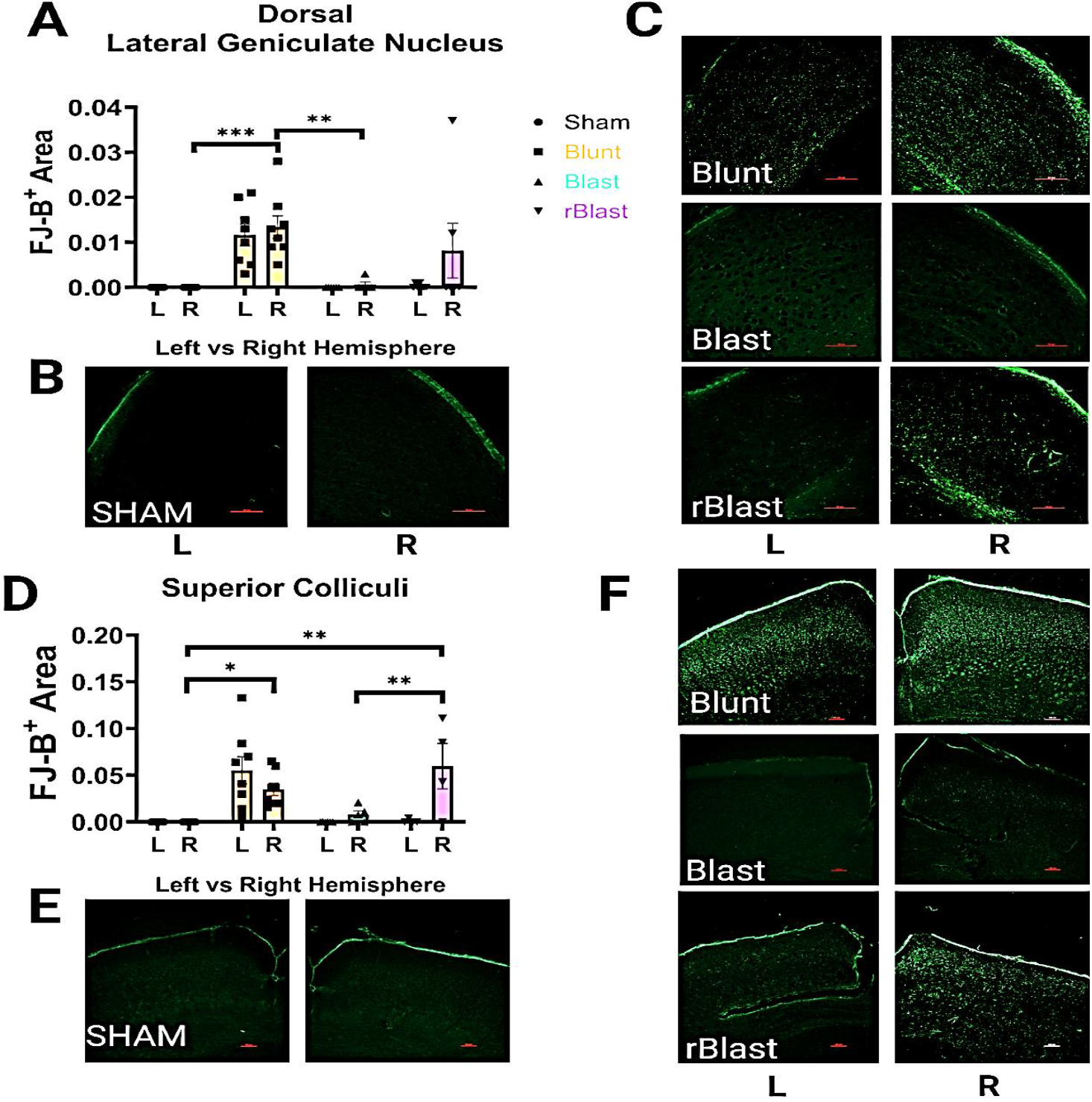
Repeat blast injury is required before degeneration is seen in OT projection regions. (A) Significant increases in FJB were only found in blunt injured mice in the dorsal LGN compared to sham, and this degeneration was also higher in blunt than blast mice. Of note, there was degeneration present in some rblast mice in the right hemisphere alone. (B) 10x representative images of sham LGN and (C) injured groups’ LGN. (D) There was also significant FJB increase in left and right blunt-injured superior colliculi compared to sham, but repeated blast injuries are required for FJB presence, which is limited to the right hemisphere. (E) 10x representative images of sham SC and (C) injured groups’ SC. * p<0.01, ** p<0.001, ***p<0.0001, scale bars represent 100μm. LGN = lateral geniculate nucleus, L= left, R=right, SC = superior colliculi, vs = versus

### Retinal cell loss is comparable in both eyes and in both models

Despite unilateral degeneration and predominantly right localized APP staining in blast mice, western blotting of retinal samples for the retinal ganglion cell (RGC) marker RBPMS revealed similar loss of RGCs in all injured mice compared to sham (p<0.001; Fig. 6). Additional examination of RGCs for markers of astrocyte reactivity (GFAP), late-stage apoptosis (PARP1), and ER stress activation (BiP; a mechanism associated with cell death in the blunt injury model)(Hetzer et al., 2021a) revealed no significant differences (data and methods described in supplementary materials; Sup Fig.2)

**Figure 6.**
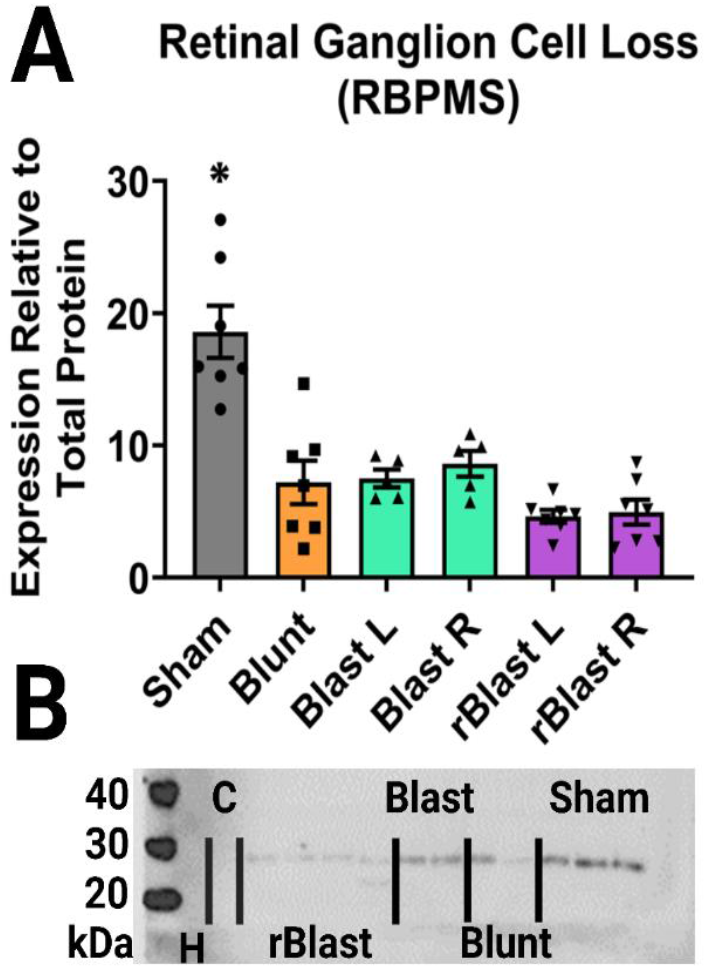
Retinal ganglion cell loss occurs in both retinas after TBI. (A) Western blot analysis for the RGC marker, RBPMS, revealed significantly reduced RBPMS in all injured retinas including both left and right for blast injured mice. (B) A representative blot (chemiluminescent) shows our negative control (H; water), intermembrane control (C; brain homogenate), and rBlast, blast, blunt, and sham lanes divided byblack lines. (C) Provides a representative image of the total protein stain used for normalization. * p<0.01. L= left, R=right, kDa = kilodalton, rBlast = repeat blast TBI

### Impaired optokinetic nystagmus is more pronounced in blunt injured mice

We examined three spatial frequency gratings to assess a range of mouse visual acuity with 0.26 cpd representing the “peak” of mouse visual acuity, 0.32 cpd being a bit thinner and harder to differentiate, and 0.39 being just below the murine visual threshold.(Hetzer et al., 2021a; Shi et al., 2018) As described in previous blunt studies, (Hetzer et al., 2021a) blunt injured mice have significantly reduced OKR responses compared to controls at each of the three cpd’s (p<0.001, p<0.001, p=0.01 respectively; Fig. 7a). Singly injured blast mice only show significant reductions at 0.32 cpd (p=0.004) and rblast mice have significantly impaired OKR at 0.26 (p=0.02) and 0.39 cpd (p=0.03). After finding that blast injury induced a predominantly unilateral, right sided, injury, we separated out OKR analyses to ask whether visual deficits would be different when the OKR response is left (i.e., clockwise rotation) vs right eye (i.e., counterclockwise rotation) dominant. There were no individual effects between groups, but there was a main effect of rotation at 0.26 cpd (p=0.02) such that the right eye had more responses than the left overall in injured groups (Fig. 7c). At 0.32 there was a main effect of injury (p<0.001) where rblast mice responded similarly to sham in the counterclockwise direction (Fig. 7d). Additionally, there was a significant subject effect at 0.39 cpd (p=0.004) such that individual rblast mice performed significantly better during counterclockwise (right eye) rotation (Fig. 7e).

**Figure 7.**
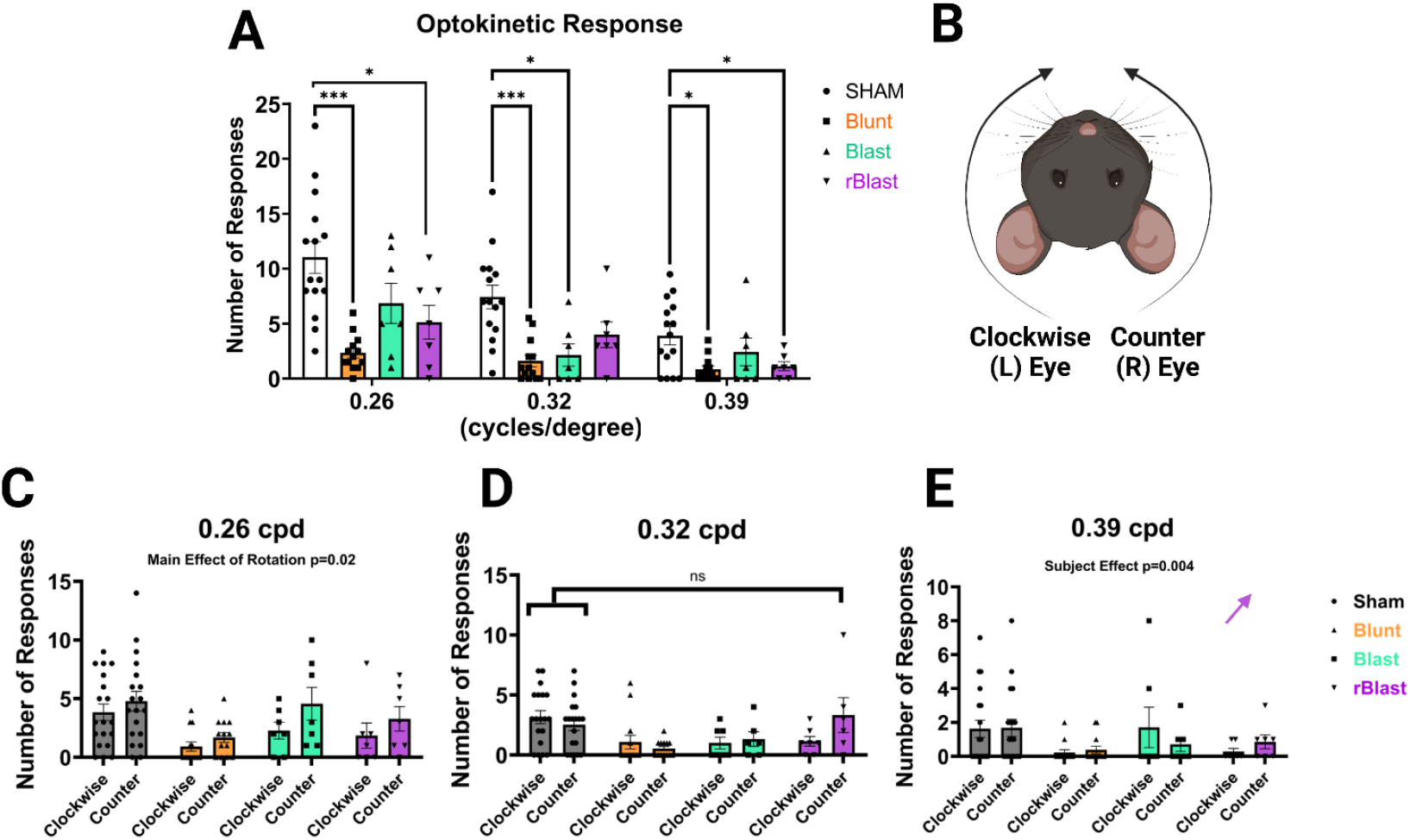
Blunt mice have more sever OKR dysfunction. (A) The number of OKR responses was tallied for each grating revealing significant blunting of this response across contrasts in blunt mice with single blast only significantly reducing OKR at 0.32, and repeat blast at 0.26, and 0.39. (B) shows an illustration of the predominant eye associated with each direction of the spinning OKR drum with the clockwise direction predominantly relying on the left eye, and vice versa. After finding unilateral OT degeneration, we analyzed our OKR data according to dominant eye rotations and found that there was a more severe left eye deficit than right at (C) 0.26cpd, no effect of direction at (D) 0.32, and significant subject differences between left and right eye dysfunction in blast injured mice at (E) 0.39 cpd. The purple arrow shows the direction of subject effects in rblast mice such that they appear to perform better with their right eye. * p<0.01, ***p<0.0001. cpd = cycles per degree, L = left, R = right, ns = not significant, rBlast = repeat blast TBI

## Discussion

We show that two different models of traumatic brain injury lead to similar deficits in vision without completely overlapping pathologies. Both blunt impact and blast-induced inertial head injuries led to comparable deficits in optokinetic responses and degeneration in the optic tract. While neurodegeneration was observed within both injury paradigms, the patterns of neurodegeneration were model- and severity-specific, which impacted both the degree and regionality of degeneration. Blunt injury led to worse bi-lateral degeneration in all vision-associated brain regions assessed, while degeneration after blast injury was limited to the optic tract and the LGN of the right hemisphere until repeated injuries were sustained. Despite unilateral degeneration in blast-injured mice, RGC loss was comparable across each method and severity indicating that an insult to one hemisphere is sufficient to induce significant loss of RGCs in both eyes, likely leading to the comparable OKR deficits observed.

Having shown that blunt injury led to significant impairment in the visually evoked optokinetic response, we asked whether blast injury would induce similar functional deficits. Our results indicate that, while there is an observed deficit in all groups, blunt injury results in more severe/pronounced deficits across the spectrum of visual gratings tested whereas blast-injured subjects demonstrated slightly better retention of OKR function at 0.26 and 0.39 frequencies. Interestingly, while functional deficits in the OKR were less severe in our blast TBI subjects than blunt-TBI subjects, they were induced following exposure to peak overpressures much higher than those required to elicit similar OKR deficits in other blast TBI studies (Guley et al., 2016b). Though the peak overpressures produced in our model are higher, our blast model has been specifically designed to generate Friedlander waveforms which occur during an improvised IED detonation, scaled to the physiologic parameters of murine subjects (Reeder et al., 2022).

Just as there are differences between blunt and blast injuries, it is important to note that there is no standardized blast TBI device. Instead, blast devices are generally made by each lab independently, creating a range of pressure wave forms, gases vs. compressed air, localization of injury, body protection, etc. It has already been noted that although studies examining OKR after blast TBI find deficits, each model has found differences in the timing of dysfunction and potential worsening or recovery of OKR performance over time (Evans et al., 2021). It is possible that discrepancies between OKR function reported here, and those generated using other blast models, result from the differences in acceleration/deceleration inertial forces generated by the blast model and containment device. Importantly, characterization of physiologic parameters and design specifications of the model, behavioral adaptations. Tissue pathology following injury induction illustrates that the blast TBI sustained following exposure to the Friedlander waveform is a mild TBI. The blast device exclusively shields the body of the mouse, leaving the head exposed to rotational movement and inducing a coup-contrecoup-like injury in conjunction with exposure to the peak blast overpressure. The heterogeneous inertial forces sustained during blast exposure are an important feature of blast-TBI pathology and are included in the Center for Disease Control and Prevention’s (CDC) categorization of distinct blast injury mechanisms with blast TBI being a component of primary, secondary, tertiary, or quaternary blast injury ((U.S.), 2003). We successfully recapitulated these effects within a preclinical setting, but now it is necessary for the field to understand how subtle differences lead to drastic changes in phenotype driven by distinct secondary injury mechanisms.

One such discrepancy becomes apparent when confirming RGC loss. Most blast models consistently report RGC layer thinning and cell loss, but some found very little (Bricker-Anthony et al., 2014), and others only examine the injured eye using the opposite eye as a within-subjects control (Allen et al., 2018; Bricker-Anthony et al., 2014). We show similar relative loss of RGCs by western blot between both models. Additional markers used to assess cell death and a potential ER-stress associated mechanism did not reveal significant differences between groups. The blunt model used has previously shown higher levels of ER stress markers up to 30 days post injury in adolescent mice, but we found no indications of ER stress in either injury group in these cohorts (Supplementary Fig. 2F). Two important differences are worth noting here. First, ER stress was measured using a less robust/direct measurement of ER stress – BiP – which we have not yet shown elevated in either model. Here we chose an ubiquitous/more global marker for ER stress due to limited tissue quantity, but it is possible that by examining other markers of ER stress, like phosphorylated eIF2α or CHOP, we might see differences. Second, that the mice tested in these experiments were adult-age mice, which may have been less severely impacted (Guilhaume-Correa et al., 2020). Alternatively, it could be that too many RGCs were lost by 7 DPI to detect any significant changes in ER stress, which could have resolved to eliminate all the injured cells by 7 days. We cannot say this conclusively, though, because we have not yet identified the time point of maximal cell loss in either of these models, and there are wide ranges reported in the TBI literature (Boehme et al., 2021; Bricker-Anthony et al., 2014; Guley et al., 2016a; Harper et al., 2019; Jha et al., 2018; Patel et al., 2017; Struebing et al., 2018).

Another possibility is that the mouse retina is predominantly composed of photoreceptors with the less numerous cones still outnumbering RGCs 4:1 (Jeon et al., 1998), leaving comparatively few RGCs even in healthy tissue. Future studies could determine RGC-specific mechanisms by sorting out RGCs. Moreover, it may be beneficial to sort astrocytes, as astrocytes have their own ER stress response (Sims et al., 2022). However, it is unlikely that astrocytes induced detectible ER stress in our samples since we also showed no increased GFAP expression in the retina (sup. fig. 2), and a reactive astrocyte phenotype is associated with ER stress upregulation in these cells (Jin et al., 2018; Levine et al., 2016; Sims et al., 2022).

We next examined brains for evidence of neurodegeneration using FluoroJade-B. Doing this, we observed a pattern of axonal degeneration consistent with previous studies in blunt injured mice with axonal degeneration seen bilaterally in subcortical visual regions (LGN, DNT, SC, OT, chiasm) (Evanson et al., 2018; Hetzer et al., 2021b). Surprisingly, however, blast tissue only exhibited unilateral FJ-B staining in the right hemisphere, which is contralateral to the blast exposure. Moreover, we only found positive staining in the optic chiasm/tract and direct optic nerve projection region, the LGN, of blast mice. It appears that, to cause degeneration in secondary or more distal projection targets, like the SC, multiple blast injuries are necessary. Other models of blast injury have shown predominantly unilateral degeneration in the optic nerve and tract (Guley et al., 2016a), but we could rarely find any reports showing this was also true in projection regions (Koliatsos et al., 2011). However, a blast model using an adapted paintball gun features glial reactivity in unilateral projection regions at high pressure (Guley et al., 2016a; Reiner et al., 2015). The easiest way to explain this is likely through injury mechanism whereby the blunt injury induces downward force over the whole optic nerve, while the blast mechanism leads to more shearing and rotational forces that might lead to worse injury unilaterally, and contralateral to the blast wave (potentially due to a contrecoup force).

Because most other blast models describing optic injury have concluded that unilateral damage/cellular response is correlated with degeneration limited to one of the optic nerves, we also sought to locate sites of axon breakage in these mice. To do this, we analyzed Amyloid Precursor Protein by immunohistochemistry. APP staining has long been employed as a measure for detecting axonal injury after TBI (Blumbergs et al., 1995; Bramlett et al., 1997; Gentleman et al., 1993; Pierce et al., 1996). APP staining revealed limited areas of damage in these models even when repeated injuries were sustained. As expected, there was no positive staining in sham mice. Unexpectedly, however, there was also almost no APP in the brains of blunt injured mice save for a few patches in the optic tracts, so we examined optic nerves and identified APP signal spanning about 1.2mm of optic nerve just proximal to the chiasm. These findings suggest that the blunt injury induces axonal breakage in the optic nerve at this location. The spread of APP signal in these blunt optic nerves is consistent with stretching of these axons as they leave the protection of the boney optic canal. We cannot definitively report this as the biomechanical mechanism, though, because compaction/crush of the nerve between the brain and skull is also possible.

In blast injured mice, APP built up in the optic tracts and ventral lateral geniculate nuclei bilaterally, which was interesting paired with unilateral FJ-B staining. Somatic FJ-B staining was not detected, but this is not uncommon in milder TBI paradigms. It is possible that APP in the LGN is not the result of primary axotomy (i.e., direct axon breakage), rather it may be evidence of stretch injury due to the small breaks of microtubules in non-degenerating cells that are still affected by injury (Hånell et al., 2015; Tang-Schomer et al., 2012; Tang-Schomer et al., 2010). It is also possible that APP served a neuroprotective function,(Hefter D. and A., 2017) at least in the left hemisphere, where there was less axonal degeneration, but future studies would need to confirm this. Future studies will also need to examine tissue both more acutely and longitudinally to determine this relationship between degeneration and APP expression as another blast model only reported APP expression in the cortex despite RGC loss and optic nerve degeneration, once again highlighting differences that must be considered even within blast models (Harper et al., 2019).

The presence of APP in the LGN could also indicate that this coup-contrecoup injury has a wider impact zone leading to broader axonal injury. This is supported when noting that APP staining was also present in a couple of regions outside the optic system in blast mice including the striatum, hypothalamus, and subventricular zones, but this was inconsistent among injured mice (data not shown). The restriction of injury to the ventral LGN also poses an intriguing finding as this region does not project to visual cortex (V1) and is not associated with “direct” visual circuits. Instead, the vLGN is more complex, including numerous inhibitory and excitatory projections between the brainstem (SC, locus coeruleus, pretectum), the accessory optic system, and the hypothalamus and predominantly responds to bright light cues (unlike its SC counterpart) perhaps explaining our less robust declines in OKR in blast mice, a response that relies on dark cues processed by the accessory optic system and SC (Ciftcioglu et al., 2020; Harrington, 1997). After noting unilateral degeneration, we also wondered if our behavioral tests were sensitive enough to detect bias in visual function. So, we separated the OKRs into those predominantly triggered by the left visual field (clockwise drum rotation) from those triggered by the right (counterclockwise rotation). These data revealed a main effect of rotation in both blast groups, which tended to have more responses during the counterclockwise rotation at 0.26 cpd. These data suggest more severe damage to the axons of the left visual field, which leave the right side of the optic chiasm to the right hemisphere of the brain. Some TBI studies have reported this bias, but generally only if the blast was directed at the eye (Allen et al., 2018). That said, left and right eye function are not often parsed apart using OKR. Further research is needed to gain an understanding of the mechanism resulting in unilateral degeneration and visual deficits despite ubiquitous RGC loss in both eyes of blast groups. An interesting extension of this research would be to determine the significance of injury severity and whether mice might be able to recover from a single versus multiple blasts.

## Conclusions

We sought to identify divergent and conserved features of injuries to the visual system following blunt and blast traumatic brain injury using murine models. In conducting these studies, we delineated the modality-specific injury of two commonly utilized preclinical TBI paradigms as they pertain to the generation of visual deficits and axonal degeneration with high spatial resolution. Specifically, we reported robust changes in the localization of optic tract degeneration between blunt and blast TBI models, with blunt neurotrauma inducing bilateral optic tract degeneration and single-and-repeat blast injury inducing unilateral optic tract degeneration contralateral to the hemisphere initially impacted by the blast front. Interestingly, we also observed paradigm-specific divergences in the degeneration of brain parenchyma. Compared to blunt neurotrauma which exclusively resulted in axonal degeneration within the optic tract, both single and repeat blast injury subjects exhibited varying degrees of axonal degeneration within anatomically distinct regions of the thalamus, with repeated blast exposure driving axonal degeneration in multiple subregions of the thalamus.

These model-dependent effects may be consequences of coup-contrecoup injury which results in sagittal plane-specific movement, axonal shearing, and neurodegeneration, as diffuse axonal injury is canonically associated with rotational head injury but not blunt neurotrauma. Elucidation of model-specific injury pathophysiology associated with blunt and blast TBI within the visual system may allow for the development of mechanism-derived therapeutic interventions for TBI, something which has not yet been resolved experimentally. Together, we believe these studies formalize and contextualize the modality-dependent effects of neurotrauma on the decay of the visual system following injury and offer insight into divergent phenotypes between paradigms that may be progressively refined for experimental or therapeutic applications.

## Abbreviations (alphabetical)

(APP): Amyloid Precursor Protein
(cpd): Cycles Per Degree
(DTN): Dorsal Terminal Nucleus
(ED): Emergency Department
(ER): Endoplasmic Reticulum
(FJ-B): Fluoro-Jade B
(GFAP): Glial Fibrillary Acidic Protein
(IED): Improvised Explosive Device
(IBA-1): Ionized Binding Adaptor Molecule 1
(LGN; ventral [vLGN]; dorsal [dLGN]): Lateral Geniculate Nucleus
(OT): Optic Tract
(OKR): Optokinetic Nystagmus Response
(PBS): Phosphate-Buffered Saline
(ROIs): Regions Of Interest
(rTBI): Repeat-Blast Traumatic Brain Injury
(RGC): Retinal Ganglion Cell
(rpm): Revolutions Per Minute
(RRT): Righting Reflex Time
(SC): Superior Colliculi
(TON): Traumatic Optic Neuropathy
(TBI): Traumatic Brain Injury

## Author Contributions

**Shelby M. Hetzer:** Data curation; Formal analysis; Investigation; Methodology; Validation; Visualization; Roles/Writing - original draft; Writing - review & editing. **Christopher O’Connell:** Formal analysis; Investigation; Methodology; Roles/Writing - original draft; Writing - review & editing. **Vivian Lallo:** Data curation; Formal analysis; Investigation; Roles/Writing - original draft; Writing - review & editing. **Mathew Robson:** Conceptualization; Funding acquisition; Investigation; Methodology; Project administration; Resources; Supervision; Validation; Visualization; Roles/Writing - original draft; Writing - review & editing. **Nathan K. Evanson:** Conceptualization; Funding acquisition; Investigation; Methodology; Project administration; Resources; Supervision; Validation; Roles/Writing - original draft; Writing - review & editing.

## Acknowledgements

This work was supported by the following: the University of Cincinnati Foundation, the University of Cincinnati Office of Research, a Cincinnati Children’s Hospital (CCHMC) Proctor Scholar Award, a CCHMC Research Innovation/Pilot Award, a Brain and Behavior Research Foundation NARSAD Young Investigator Award (#25230, MJR), a PhRMA Foundation Research Starter Grant (MJR), a National Institutes of Health NRSA T32, and a National Institute of Neurologic Disorders and Stroke (#NS007453).

---

**Supplementary Figure 1.**
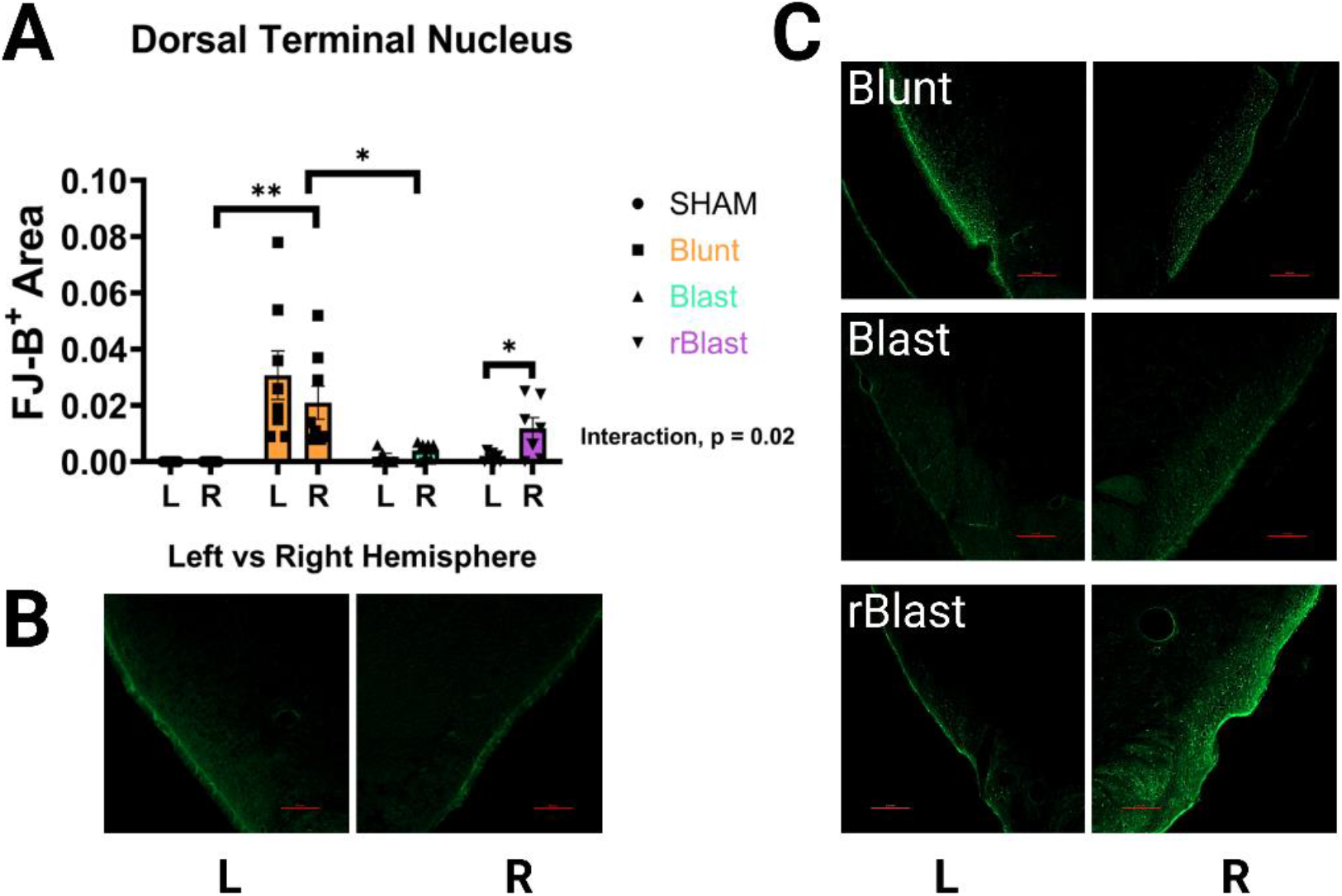
Unilateral degeneration is found in optic nerve projections in blast and blunt mice. In addition to the optic regions discussed in the main text, we also looked at the dorsal terminal nucleus, a branch of the optic nerve in the accessory optic system, which plays a role in the brain’s coordination of the optokinetic response. As with the SC, (A) we found significantly increased bilateral FJB (degeneration) in blunt injured mice (previously reported in this model (Hetzer et al., 2021b)) with only unilateral increases in rblast mice in the right hemisphere. (B) There was no positive staining in any sham mice, but positive FJB is clearly seen in (C) all injured groups including single blast, but it was not enough to reach significance. L= left, R=right, * p<0.01, ** p<0.001, scale bars represent 100 μm.

**Supplementary Figure 2.**
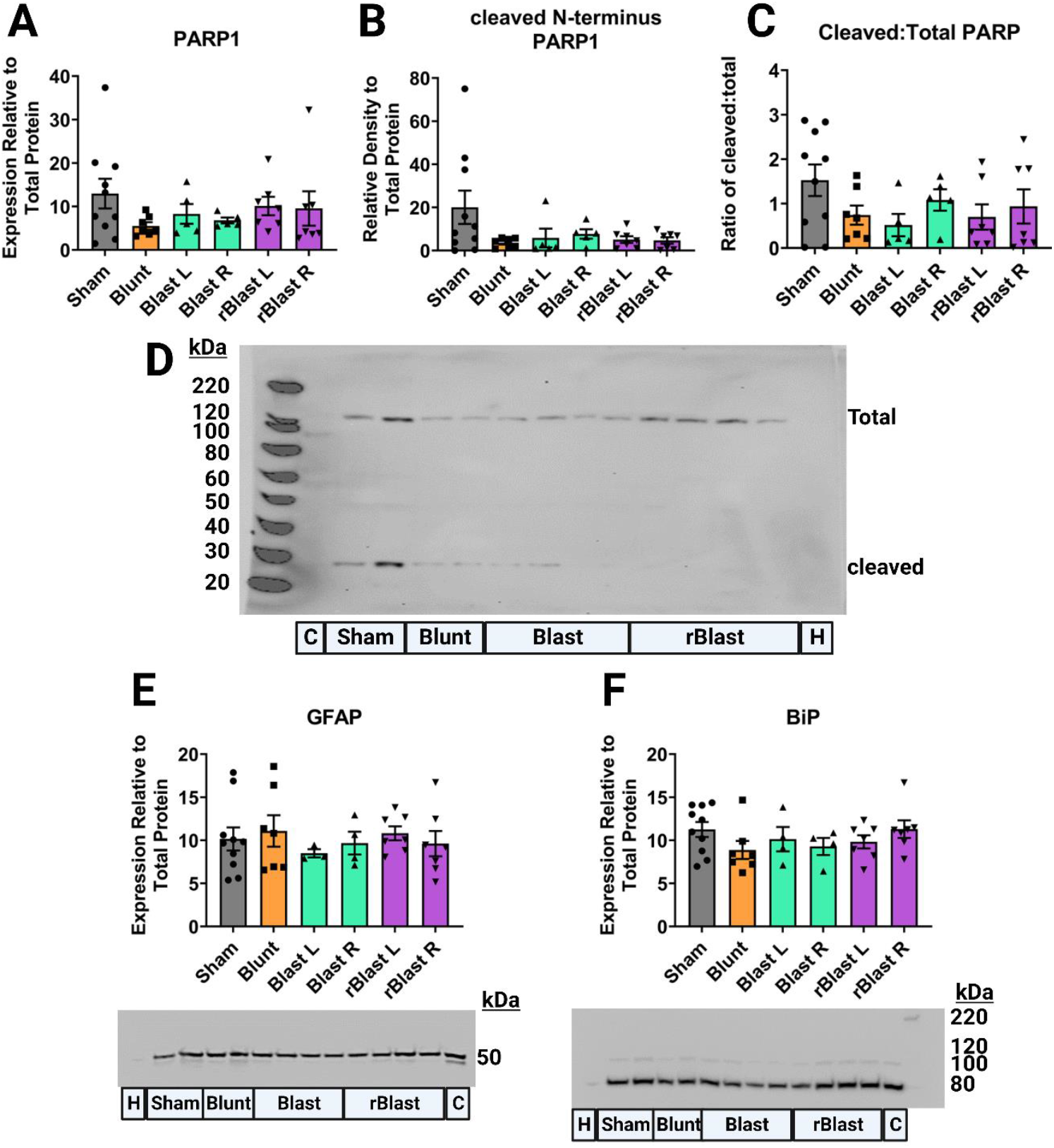
No active apoptosis, astrogliosis, or ER stress was detected 7 days post injury. We also examined retinal extracts for the late-stage apoptosis marker PARP1 but found no significant changes in (A) total PARP1, (B) cleaved N-terminal PARP1, or (C) the ratio of the two indicating that at this time point there was no active apoptosis going on. (D) shows a representative blow with the molecular weights detailed along the right side for total (∼120kDa) and cleaved-terminal PARP1 (∼28 kDa). We also wondered whether there would be increased astrogliosis in the retina after TBI but found no changes in glial fibrillary astrocytic protein (GFAP; ∼50kDa). Previous work has implicated ER stress as a mechanism of cell death that is active up to 30 days post injury, but the chosen marker for these studies, BiP (Binding Immunoglobin Protein; ∼80 kDa) was not elevated and future assays for other ER stress related proteins need to be done. kDa= kilodalton, C=intermembrane control brain homogenate, H=water negative control.

